# Combined deletion of ZFP36L1 and ZFP36L2 drives superior cytokine production in T cells at the cost of cell fitness

**DOI:** 10.1101/2024.12.11.627889

**Authors:** Nordin D. Zandhuis, Antonia Bradarić, Carmen van der Zwaan, Arie J. Hoogendijk, Branka Popović, Monika C. Wolkers

## Abstract

A key feature of cytotoxic CD8^+^ T cells for eliminating pathogens and malignant cells is their capacity to produce pro-inflammatory cytokines, which includes TNF and IFNγ. Provided that these cytokines are highly toxic, a tight control of their production is imperative. RNA-binding proteins (RBPs) are essential for the fine-tuning of cytokine production. The role of the RBP ZFP36L1 and its sister protein ZFP36L2 herein has been established, however, their relative contribution to cytokine production is not well known. We here compared the effect of ZFP36L1 and ZFP36L2 single and double deficiency in murine effector CD8^+^ T cells. Whereas single deficient T cells significantly increased cytokine production, double deficiency completely unleashed the cytokine production. Not only the TNF production was substantially prolonged in double-deficient T cells. Also, the production of IFNγ reached unprecedented levels with >90% IFNγ–producing T cells compared to 3% in WT T cells, even after 3 days of continuous activation. This continuous cytokine production by double-deficient T cells was also observed in tumor-infiltrating lymphocytes in vivo, however, with no effect on tumor growth. Rather, ZFP36L1 and ZFP36L2 double deficiency resulted in decreased cell viability, impaired STAT5 signaling, and dysregulated cell cycle progression. In conclusion, while combined deletion in ZFP36L1 and ZFP36L2 can drive continuous cytokine production even under chronic activation, safeguards are in place to counteract such super-cytokine producers.

## INTRODUCTION

Cytotoxic CD8^+^ T cells protect us from microbial infections and can fight cancer cells. Fundamental to mounting effective T cell responses is their production of pro-inflammatory cytokines (1–4). In addition, memory T cells contain ready-to deploy mRNAs, which enables them to rapidly produce cytokines upon reactivation, and thus protect us from recurring infections (5–8). However, aberrant production of these toxic cytokines can have detrimental effects, as is for instance observed upon CAR T cell therapy and in patients that severely suffer from COVID-19 infections (9, 10). Several regulatory mechanisms are therefore in place to tightly regulate cytokine production, such as transcriptional, epigenetic, and post-transcriptional regulation (11, 12).

RNA binding proteins (RBPs) are key mediators of post-transcriptional regulation. Amongst other processes, they define RNA splicing, nuclear export, translation, and stability of the target mRNA (13), with important functional consequences. For instance, the ZFP36 family proteins ZFP36, ZFP36L1, and ZFP36L2 have been shown to modulate T cell development, differentiation and function (5, 14–19). They do so by interacting with AU-rich elements (AREs) present in the 3’UTR of their target mRNAs, including cytokines (20). Interestingly, even though all ZFP36 proteins bind to the same sequence, they act in a context-dependent manner. For instance, whereas ZFP36L2 blocks translation of preformed *Tnf* and *Ifng* mRNA in resting memory CD8^+^ T cells (5), it destabilizes *Ifng* mRNA *in* continuously activated effector CD8^+^ T cells (19). Likewise, ZFP36 and ZFP36L1 can control translation of *Tnf* and *Il2* mRNA in CD4^+^ T cells (21, 22), yet in CD8^+^ T cells, the *IL2* mRNA becomes destabilized (14, 18, 19). Intriguingly, ZFP36L1-deficient tumor-infiltrating T cells (TILs) produced more IFNγ, which resulted in a modest yet significant delay of tumor outgrowth (18). ZFP36L2-deficient TILs also showed increased IFNγ production, however, they could not restrict tumor outgrowth (19). Whether ZFP36L1 and ZFP36L2 are redundant or collaborate in regulating IFNγ, and whether dual deficiency generates superior T cell responses against tumors is, however, not known.

To decipher how ZFP36L1 and ZFP36L2 regulate cytokine production in effector T cells, we determined their expression and regulation kinetics. We found that the protein expression dynamics mirrored that of their activity. Specifically, ZFP36L1 dampens cytokine expression during early T cell activation and at its peak of production, i.e. at 6h, whereas ZFP36L2 is active at later time points. Intriguingly, ZFP36L1 and ZFP36L2 double deficiency further increased cytokine expression at earlier time points. When exposed to continuous antigen, the double deficiency of ZFP36L1 and ZFP36L2 completely unleashed the production of IFNγ. High IFNγ production was also found *in vivo* in TILs, however, ZFP36L1 and ZFP36L2 double deficient CD8^+^ T cells could not control tumor growth. Rather, they showed limited cell fitness due to decreased viability and a perturbed cell cycle progression. We therefore conclude that RBPs regulate multiple modules in concert to define the fate of T cells.

## RESULTS

### Differential expression kinetics of ZFP36 family members upon T cell activation

To determine the expression of ZFP36 family members during T cell activation, we first mined the proteomics data of ImmPRes from naive OT-I T cells that were activated for up to 24h with the OVA_257-264_ peptide (23). Whereas ZFP36 expression was below the detection limit, the expression of ZFP36L1 rapidly increased upon T cell activation, peaking at 3-4h of T cell activation and declining slowly thereafter (**Fig. 1A**). ZFP36L2 expression remained low during the first 12h of activation and increased from 18h of activation onwards (**Fig. 1A**). The expression kinetics of ZFP36 proteins of *in vitro* generated effector T cells were similar. ZFP36L2 expression slowly declined at 6h of activation with OVA peptide, but it restored and increased at 24-48h of activation (**Supp. Fig 1A**). Conversely, ZFP36 and ZFP36L1 expression increased at 6h, and decreased at 24h and 48h of T cell activation (**Supp. Fig 1A**). Of note, we detected no effects on ZFP36 and ZFP36L1 protein expression in ZFP36L2^CKO^ T cells (**Supp. Fig 1A**), suggesting that compensatory effects on ZFP36 and ZFP36L1 in ZFP36L2^CKO^ T cells are limited.

**Figure 1.**
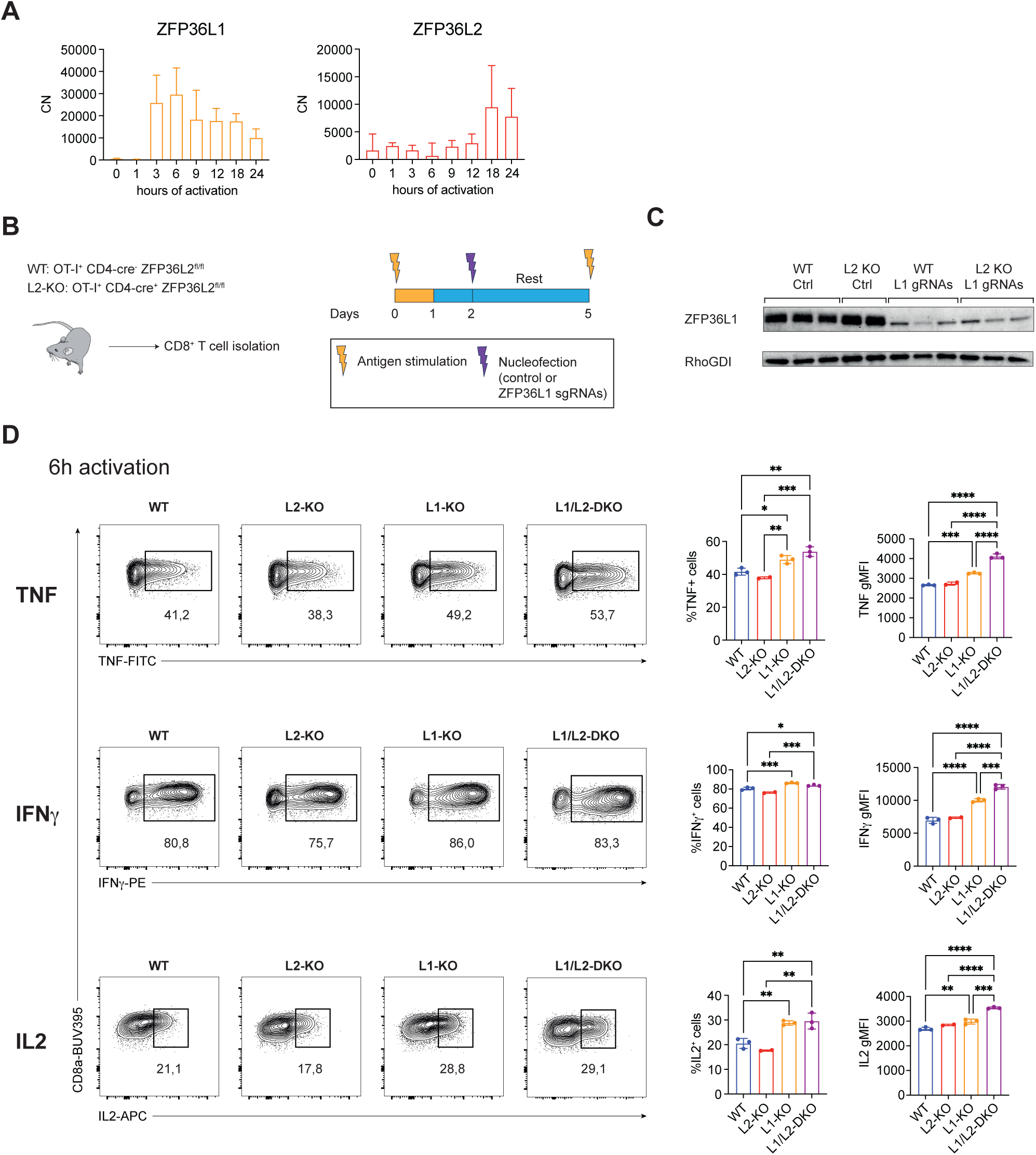
Differential expression kinetics of ZFP36L1 and ZFP36L2 during T cell activation correlates with ZFP36L1-mediated regulation of early cytokine production. (**A**) ZFP36L1 and ZFP36L2 expression in Copy Numbers (CN) from naive OT-I T cells stimulated with OVA peptide for up to 24h. Data extracted from the ImmPRes Immunological Proteome Resource (23). **(B)** Schematic overview of generation of WT, L2-KO, L1-KO and L1/L2-DKO T cells using ZPF36L2^cko^ and WT OT-I T cells treated with control sgRNA (WT, L2-KO) or ZFP36L1-targeting sgRNAs (L1-KO, L1/L2-DKO). **(C)** Immunoblot displaying ZFP36L1 protein expression in indicated T cells. RhoGDI served as a loading control. n=2-3 mice per condition. **(D)** Representative frequency of TNF, IFNγ and IL-2 producing T cells after 6h of co-culture of WT, L2-KO, L1-KO and L1/L2-DKO OT-I T cells with B16-OVA target cells. Brefeldin A was added for the last 2h of activation. Expression levels for TNF, IFNγ and IL-2 among cytokine-producing cells are displayed using geometric mean fluorescence intensity (gMFI). Each dot indicates one mouse, n=2-3 mice per condition. (A-D) Presented data is from individual experiments. (A-C) One independent experiment has been performed. (D) Data is representative for four independent experiments. Data were analyzed by one-way ANOVA with Tukey multiple comparison correction, mean ± SD (D); *P < 0.05, **P < 0.01, ***P < 0.001, ****P < 0.0001.

### ZFP36L1 controls cytokine production at early time points

Provided their differential expression kinetics, we questioned what the relative contribution of ZFP36 proteins is on cytokine expression. To study this, we compared WT, ZFP36L1- and ZFP36L2-single deficient T cells with ZFP36L1/ZFP36L2-double deficient T cells. CD4-Cre induced ZFP36L1/ZFP36L2 double deficiency results in impaired T cell priming (16), which impedes the use of this model for studies in effector T cells. To overcome this, we gene-edited WT and ZFP26L2^CKO^ OT-I T cells with ZFP36L1-targeting CRISPR Cas9 gRNAs, or with non-targeting controls to obtain WT T cells, ZFP36L1 (L1-KO) and ZFP36L2 (L2-KO) single and double deficient (L1/L2-DKO) T cells (**Fig. 1B, C**).

We next measured the cytokine production of T cells with the differential RBP deletion upon exposure to B16-OVA target cells (24). In concordance with their respective expression kinetics (**Fig. 1A**), the production of TNF, IFNγ and IL-2 was significantly increased at 6h of activation in L1-KO cells compared to WT T cells, but not in L2-KO T cells (**Fig. 1D**). Interestingly, double deficient L1/L2-DKO T cells displayed higher percentages of cytokine-producing T cells than L1-KO cells and cytokine production per cell, as defined by the Geometric Mean Fluorescence intensity (GeoMFI; **Fig. 1D**). Thus, ZFP36L1 primarily regulates the cytokine production of T cells at early time points. ZFP36L2 also contributes, but only in the absence of ZFP36L1.

### Double deficiency of ZFP36L1 and ZFP36L2 unleashes cytokine production

We previously showed that ZFP36L2 dampens the production of IFNγ when T cells are continuously exposed to antigen (19). To test how L1-KO and L1/L2 DKO T cells responded to continuous antigen exposure, we re-exposed effector T cells each day to freshly seeded B16-OVA cells for a total period of 96h (**Fig. 2A**). To measure cytokine production, we added Brefeldin A for the last two hours of activation. As we previously showed (6, 18, 25), the production of TNF and IL-2 was lost in WT T cells after 24h of T cell activation (**Supp. Fig. 2A, B**). Whereas ZFP36L1/L2 single and double deficiency did not alter the kinetics of IL-2 production at 24h (**Supp. Fig. 2A**), this was not the case for TNF. TNF production was still detectable in L2-KO T cells, and even more so in L1/L2-DKO T cells (**Supp. Fig. 2B**). Even at 48h and 72h of continuous activation, TNF production was with ±23% at 48h, and with ±10% at 72h still measurable in L1/L2 DKO T cells **(Supp. Fig. 2B**). Only at 96h, the TNF production was lost in all genetic backgrounds **(Supp. Fig. 2B**).

**Figure 2.**
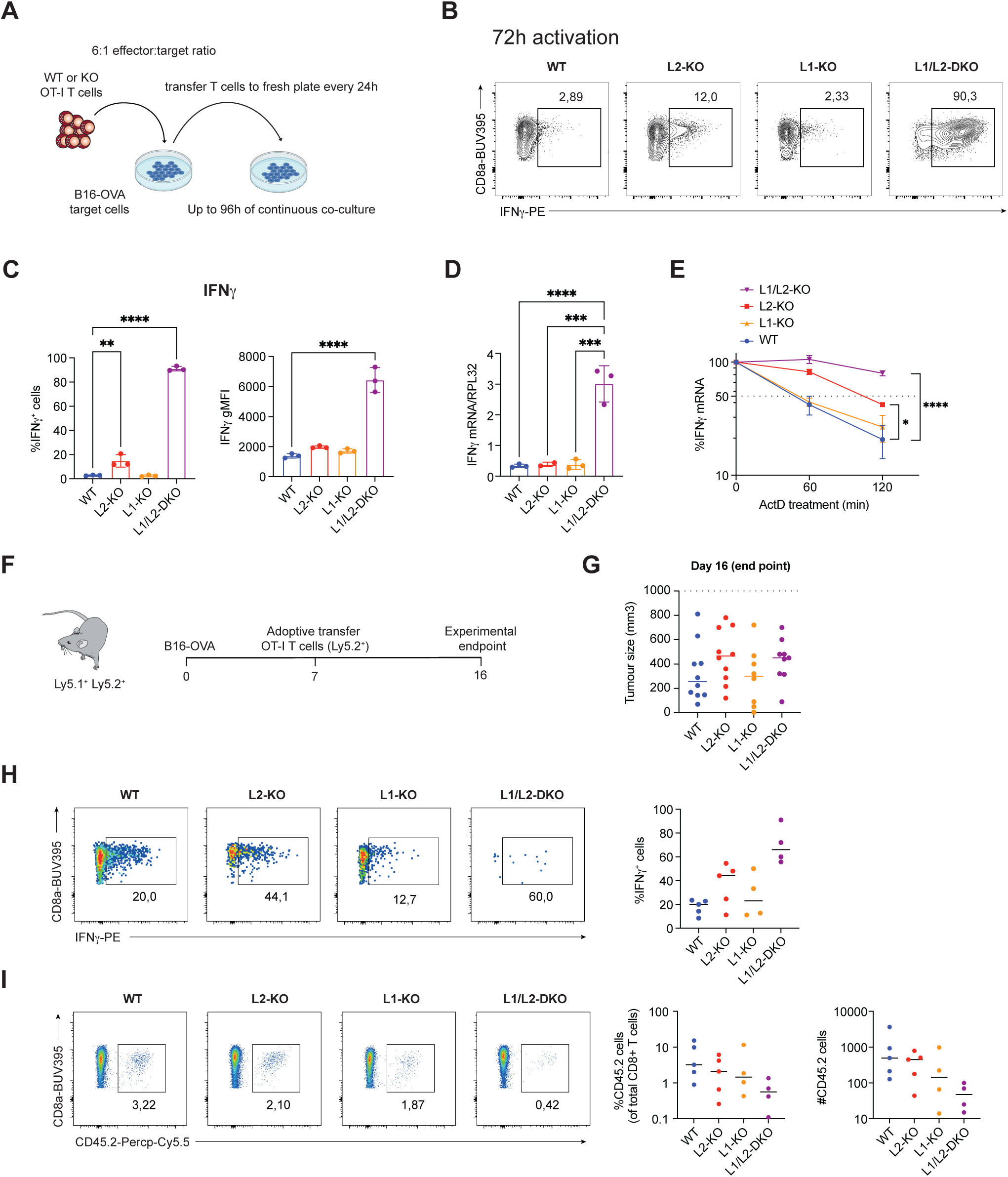
ZFP36L2+ZFP36L1 double deficiency results in superior IFNγ production during continuous activation, but at the expense of cell loss. **(A)** Schematic representation of the *in vitro* B16-OVA co-culture system to continuously stimulate OT-I T cells for up to 96h. Every 24h, T cells were transferred to a fresh plate of pre-seeded B16-OVA cells. **(B)** Representative production of IFNγ by WT, L2-KO, L1-KO and L1/L2-DKO T cells upon 72h of continuous exposure to B16-OVA cells. Brefeldin A was added for the last 2h of activation. **(C)** Frequency of IFNγ producing T cells (left panel) and gMFI levels of IFNγ (right panel) among IFNγ producing cells after 72h of continuous exposure to B16-OVA cells. n=3 mice per condition. **(D)** *Ifng* mRNA levels and (**E**) *Ifng* mRNA decay normalized to *Rpl32* mRNA levels in OT-I T cells after 72h of continuous co-culture with B16-OVA. For mRNA decay, OT-I T cells were treated with 1μg/ml Actinomycin D (ActD) for indicated time points. n=2-3 mice per condition. **(F)** Schematic overview of *in vivo* B16-OVA tumor model. **(G)** Tumor volume at day 16 post tumor inoculation of mice that received indicated OT-I T cells. n=10 for WT mice, n=10 for L2-KO mice, n=8 for L1-KO mice and n=9 for L1/L2-DKO mice. **(H)** Frequency of IFNγ producing T cells among CD45.2^+^ tumor-infiltrating OT-I T cells after 4h of *ex vivo* culture in the presence of Brefeldin A and Monensin. **(I)** Frequency and absolute number of CD45.2^+^ OT-I T cells isolated from B16-OVA tumors. (H,I) n=4-5 mice per condition. (C-E, G-I) Presented data is from individual experiments. (C) Data is representative for three independent experiments. (D-E, G-I) One independent experiment has been performed. Data were analyzed by one-way ANOVA with Tukey multiple comparison correction, mean ± SD (C, D, E); *P < 0.05, **P < 0.01, ***P < 0.001, ****P < 0.0001.

We then turned our attention to IFNγ. As expected, substantial IFNγ production was measured in WT T cells at 24h and 48h of activation (**Supp. Fig. 2C)**. In concordance with its expression kinetics, L2-KO T cells produced more IFNγ at 24h and 48h of continuous T cell activation than WT or L1-KO T cells (**Supp. Fig. 2C)**. Strikingly, whereas WT and L1-KO T cells ceased to produce IFNγ at 72h, almost all L1/L2-DKO T cells maintained their production of IFNγ, both in terms of percentage and production per cell (**Fig 2B, C**). Even at 96h of continuous T cell activation, the production of IFNγ was with 32.4 ± 3.5% still significant in L1/L2-DKO T cells (**Supp. Fig. 2C**). L1/L2-DKO T cells contained 10-fold higher *Ifng* mRNA levels, which was at least in part due to an almost complete loss of mRNA decay, as determined by blocking *de novo* transcription with Actinomycin D (ActD) (**Fig. 2D, E**). In conclusion, ZFP36L2 dominates the regulation of cytokine production at later time points. Yet, double depletion of ZFP36L1 and ZFP36L2 unleashes the IFNγ production during continuous T cell activation.

### Loss of tumor-infiltrating ZFP36L1 and ZFP36L2 DKO T cells *in vivo*

IFNγ is critical for effective T cell responses against tumors (1, 26). Based on their superior IFNγ production upon continuous T cell activation *in vitro*, we hypothesized that this feature could render L1/L2 DKO T cells better tumor suppressors than WT T cells. To test this, we adoptively transferred OT-I T cells with the four different genetic backgrounds into B16-OVA tumor-bearing mice. 9 days later, we measured tumor size and the production of IFNγ in tumor-infiltrating T cells (TILs) upon 4h of incubation with Brefeldin A and Monensin (**Fig. 2F**). Surprisingly, we observed no significant changes in tumor size (**Fig. 2G**). Even though the production of IFNγ appeared higher in L1/L2 DKO TILs (**Fig. 2H**), the yield of transferred L1/L2 DKO TILs was substantially lower than that of any other genetic background (**Fig. 2I**). Thus, even though ZFP36L1 and ZFP36L1 double deficiency relieves the block on IFNγ production within tumors, this occurs at the cost of cell loss.

### Protein expression of different gene sets is altered in L1/L2-DKO T cells

We next aimed to uncover why L1/L2-DKO T cells lost their cell fitness *in vivo*. Albeit less prominent, lower cell viability in L1/L2-DKO T cells was also observed in *in vitro* cultures, both in resting and reactivated T cells (**Fig. 3A, B**). We therefore turned to the *in vitro* model to compare the protein expression of resting and activated T cells of all four genetic backgrounds by Mass Spectrometry. Of the 8040 identified proteins, 678 proteins in resting and 160 proteins activated T cells were found to be statistically significant differentially abundant (adjusted P-value < 0.05 and log2 fold change < or > 0.5, indicated throughout with an asterisk, **Supp. Table 1**) between WT and L1/L2-DKO T cells (Fig 3C). As expected, the ZFP36L1 and ZFP36L2 protein levels were substantially lower in the respective knockout T cells, and effects on ZFP36 and ZFP36L1 in L2-KO and L1/L2-DKO cells were limited (**Supp. Fig 3A**).

**Figure 3.**
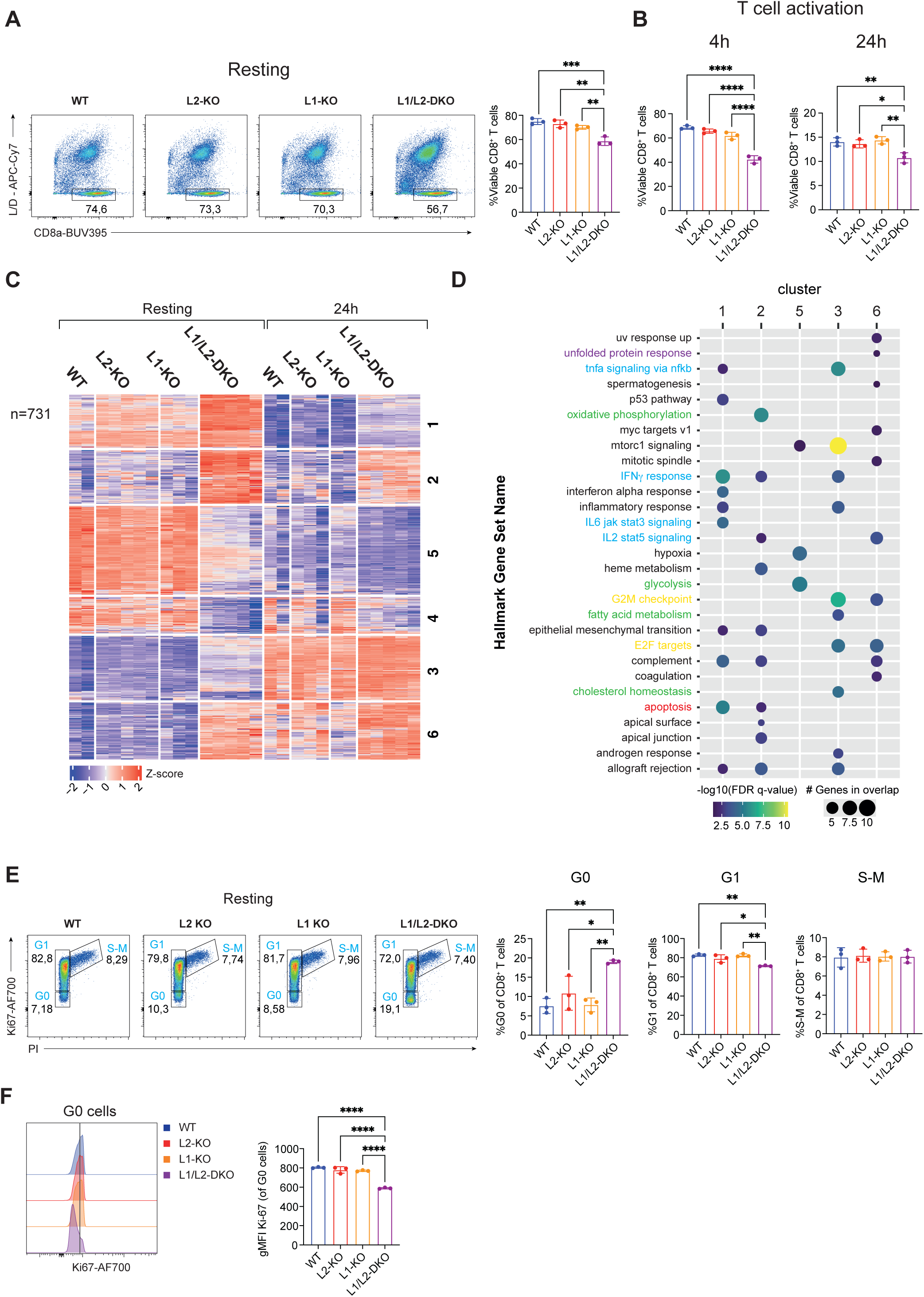
Decreased cell viability of L1/L2-DKO T cells, which show a dysregulated cell cycle progression. (**A-B**) Frequency of viable (LiveDead^negative^) T cells among resting (A) OT-I T cells (4 days after initial activation), and **(B)** after 4h and 24h of activation with the OVA peptide. n=3 mice per condition. **(C)** Heatmap of Mass spectrometry analysis showing unsupervised clustering of differentially expressed proteins (P.adj < 0.05 & Log2FC > or < 0.5) between WT and L1/L2-DKO cells of resting T cells and after 24h of activation. Number of clusters was determined using k-means clustering. n=2-5 mice per condition. **(D)** Gene Set Enrichment Analysis (GSEA) for each cluster identified in (C). Gene sets associated with unfolded protein response (purple), cytokine signaling (blue), metabolism (green), cell cycle progression (yellow) and apoptosis (red) are highlighted. **(E)** Cell cycle analysis of resting OT-I T cells using the proliferation marker Ki-67 and propidium iodide (PI). Left: representative dot plots. Right: compiled data of percentages of G0, G1 and S-M phase. n=3 mice. **(F)** Expression levels of Ki67 among G0 (Ki67^low^ PI^negative^) cells. (A-F) Presented data is from individual experiments. (A) Data is representative for three independent experiments. (B-D) One independent experiment has been performed. (E-F) Data is representative for two independent experiments. Data were analyzed by one-way ANOVA with Tukey multiple comparison correction, mean ± SD (A, B, E, F), *P < 0.05, **P < 0.01, ***P < 0.001, ****P < 0.0001.

To determine which proteins were direct targets of ZFP36L1, we re-analyzed ZFP36L1 individual-nucleotide resolution UV crosslinking and immunoprecipitation (CLIP) data from activated WT and L1-KO T cells (14). Of the 1965 regions we identified in 521 transcripts, 439 interactions were located in intronic regions, known to contain AREs (27) (**Supp. Fig. 3B-C)**. 60 interactions were located in the 3’UTR of target genes, including *Tnf*, *Ifng,* and *Il2* mRNA (**Supp. Fig. 3C-D**). Indeed, the ZFP36L1 iCLIP targets genes *Lta, Pim1, Ifng* and *Cdkn1a* also showed higher protein intensity values in the MS analysis of resting L1/L2-DKO T cells (**Supp. Fig. 3E, F**). After 24h of activation, ZFP36L1 iCLIP target genes *Tnf, Cdkn1a,* showed significantly higher protein levels, and *Ccl3, Ccl4* and *Il3* a trend of higher protein intensity levels in L1/L2-DKO T cells (**Supp. Fig. 3E, F**). Intriguingly, CCL3 protein expression was below detection limit in resting WT and most single KO T cells, yet measurable in L1/L2-DKO T cells **(Supp. Fig 3F**), indicating that double deficiency can unleash protein expression of certain target genes.

Clustering analysis of proteins with different intensity values between WT and L1/L2-DKO T cells revealed 6 gene clusters (**Fig. 3C**). Gene Set Enrichment Analysis (GSEA) using Hallmark gene sets of these clusters uncovered that in L1/L2-DKO T cells, proteins associated with cytokine signaling (TNF signaling via NFKB, IFNγ response, IL-2-STAT5 signaling), metabolism (oxidative phosphorylation, glycolysis and fatty acid metabolism), cell cycle progression (G2M checkpoint, E2F targets), unfolded protein response (UPR), and apoptosis are enriched (**Fig. 3D**). While we only found three proteins associated with the UPR to be differentially abundant in resting L1/L2 DKO T cells, the key ER-stress sensor PERK (Eif2ak3) was significantly increased in resting and activated L1/L2-DKO T cells, indicating a potentially increased UPR response (**Supp. Fig. 3G, H**).

### Cell cycle progression is altered in the absence of ZFP361 and ZFP36L2

Proteins involved in the cell cycle progression (E2F targets and G2M checkpoint) included the key cell cycle regulatory genes Cks2, Cdc6, Ccne1 and Cdkn1a, and were almost exclusively upregulated in resting L1/L2-DKO T cells (**Supp. Fig, 3I, J**). When measuring the cell cycle in resting WT, L1-KO and L2-KO T cells, we found similar frequencies of T cells in the G0, G1, and S-M phase (**Fig. 3E**). However, L1/L2-DKO T cells had slightly decreased percentages of cells in the G1 phase, and notably, 2-fold more cells in the G0 phase (**Fig. 3E**). The expression level of Ki-67 in the G0 phase of L1/L2-DKO T cells was also substantially lower, which, according to the relationship between Ki-67 expression and duration of cells spent in quiescence (28), indicates that L1/L2-DKO T cells remained for a longer time in quiescence (**Fig. 3F**). Furthermore, L1/L2-DKO T cells were dying more when in the G0 and S-M phase than T cells from the other genotypes (**Supp. Fig. 4A**). The block in G0 was still observed when L1/L2-DKO T cells were activated with OVA peptide for 4h (**Supp. Fig. 4B**). After 24h of activation, however, T cells started to divide, and the frequency of G0 cells was similar between WT and L1/L2-DKO T cells (**Supp Fig. 4C**). Indeed, L1/L2-DKO proliferation was similar or even slightly increased compared to the other genotypes (**Supp. Fig. 4D**). Thus, ZFP36L1/ZFP36L2 double deficiency attenuates the cell cycle progression, which is however relieved upon T cell activation. Together, these data show that the induction of cell cycle quiescence in resting cells is partially regulated by the RBPs ZFP36L1 and ZFP36L2.

**Figure 4.**
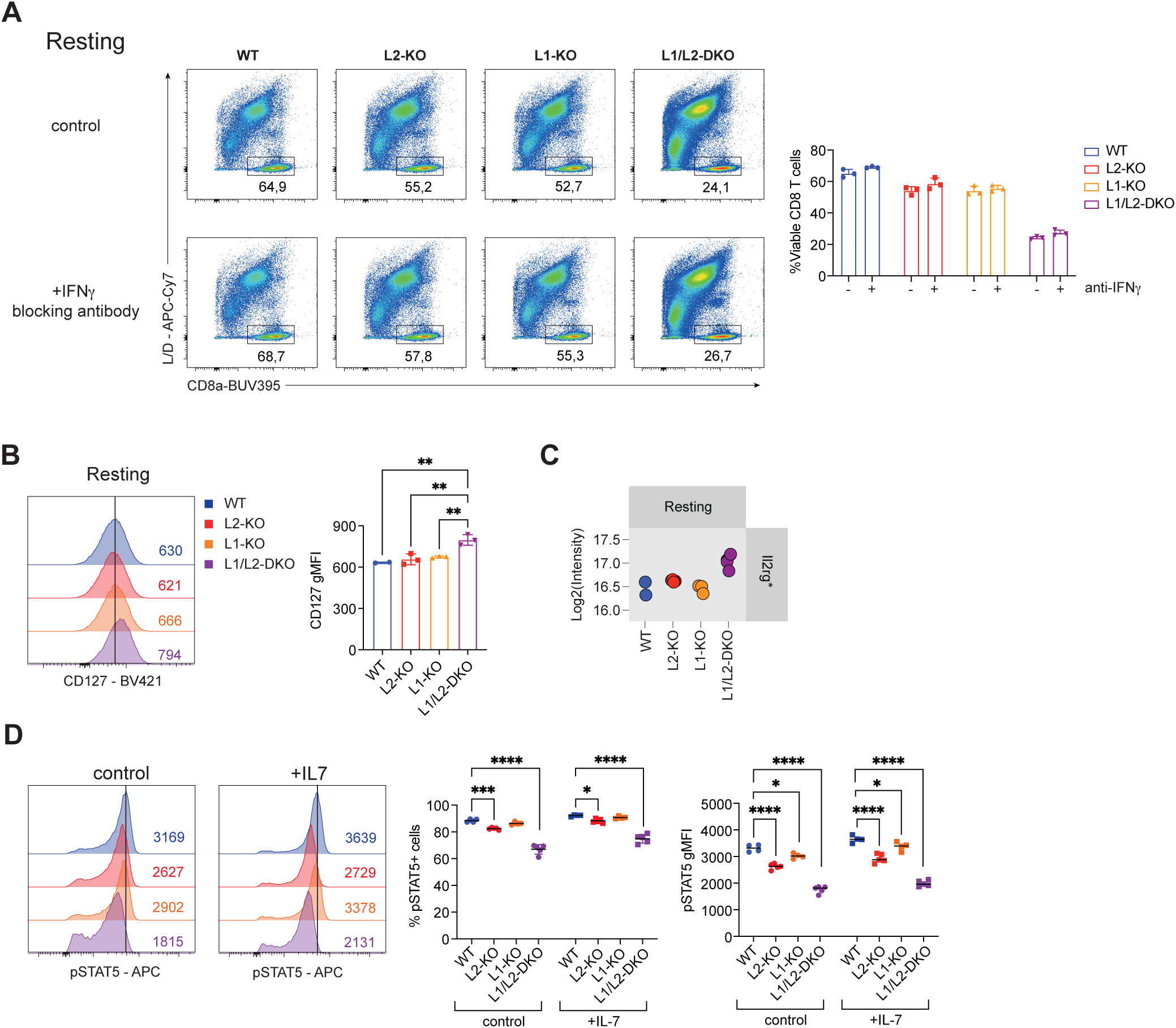
L1/L2-DKO T cells T cells show impaired IL-7 signalin. **(A)** Left: Representative analysis of frequency of viable (LiveDead^negative^ CD8^+^) T cells among resting OT-I T cells in the absence or presence (10μg/ml) of an IFNγ blocking antibody. Right: data compiled from n=3 mice. **(B)** Expression levels of IL-7Ra (CD127) among resting T cells displayed using gMFI. n=3 mice. **(C)** Protein expression levels in log2 intensity for IL-2Rg in resting T cells. n=2-5 mice. **(D)** Frequency of phosphorylated (Tyr694) STAT5 (pSTAT5) positive cells and pSTAT5 levels as defined by gMFI levels among resting T cells. n=4-5 mice. (A-D) Presented data is from individual experiments. (A-D) One independent experiment has been performed. Data were analyzed by two-sided, paired Student’s t-test, mean ± SD (A), or by one-way ANOVA with Tukey multiple comparison correction, mean ± SD (B, C, D); *P < 0.05, **P < 0.01, ***P < 0.001, ****P < 0.0001. Asteriks (C) indicates significantly differentially expressed proteins between WT and L1/L2-DKO cells; P.adj < 0.05 & Log2FC > or < 0.5.

### IL-7R signaling is perturbed in the absence of ZFP36L1 and ZFP36L2

We also found and enrichment of proteins involved in the IFNγ response **(Supp. Fig. 4E)**, which were in line with the increased expression of IFNγ and IFNGR1 in L1/L2-DKO T cells (**Supp. Fig. 3F, 4F**). IFNγ can both block cell cycle progression and was suggested to promote apoptosis (29). However, adding a blocking antibody against IFNγ in T cell cultures did not rescue L1/L2-DKO T cells (**Fig. 4A**), indicating that the IFNγR signaling pathway is not responsible for the reduced cell fitness of L1/L2-DKO T cells.

Signals received through the common gamma chain cytokine receptors IL-2R, IL-4R, IL-7R and IL-15R are crucial for T cell survival (30). Interestingly, thymic CD4-Cre-induced L1/L2- DKO T cells express less CD127 (IL-7ra) and display reduced STAT5 phosphorylation, which leads to increased apoptosis (15). Because genes involved in IL-2 STAT5 signaling were enriched in L1/L2-DKO T cells compared to WT and single KO T cells (**Supp. Fig. 4G**), we questioned whether STAT5 signaling was altered in L1/L2-DKO T cells. In resting mature L1/L2-DKO T cells, the two IL-7R receptor subunits CD127 and IL-2RG were upregulated (**Fig. 4B, C**). When T cells were deprived of rIL-7 for 4 hours and then re-exposed to the cytokine for 20 min, the frequency and levels of pSTAT5 were increased across all genotypes, however to different degrees (**Fig. 4F**). In L1-KO cells, only the geoMFI was slightly lower than that of WT T cells, but not the frequency of pSTAT5 positive cells (**Fig. 4D**). L2-KO T cells had a lower frequency and levels of pSTAT5 than WT T cells (**Fig. 4D**). However, in L1/L2-DKO T cells, IL-7 signaling was most severely affected (**Fig. 4D**). In line with this finding, the phosphatase Dusp4 which dephosphorylates Stat5 (31) was slightly upregulated in L1/L2-DKO T cells (**Supp. Fig. 4H**). Of note, total Stat5a and Stat5b protein expression remained unchanged (**Supp. Fig. 4I**). In conclusion, these data show that IL-7 signaling in effector T cells is regulated by ZFP36L2 and ZFP36L1.

## DISCUSSION

In this study, we show that L1/L2 double deficiency almost completely overcomes the attenuated cytokine production upon continuous T cell stimulation. While extended production of TNF and IFNγ could be observed in L1/L2-DKO T cells, IL-2 production was undetectable after 24h of activation. Indeed, IL-2 depends more on transcriptional regulation than TNF, IFNγ, CCL3, and CCL4, which rely more on post-transcriptional regulation (PTR) for their expression (5, 6) Interestingly, even though ZFP36L2 deficiency on its own had no effects for early cytokine production (5, 19), ZFP36L1 and ZFP36L2 double deficiency showed highest cytokine expression already at this time point. This finding suggests that ZFP36L2 is outcompeted by ZFP36L1 at early time points, either due to its lower expression levels early during T cell activation, or due to lower affinity to its target mRNAs.

Irrespective of the high cytokine production, adoptive transfer of L1/L2-DKO T cells could not block tumor outgrowth. *In vitro*, cell viability could be improved by TCR triggering, but in vivo, this did not rescue the reduced cell numbers of L1/L2-DKO T cells within tumours. We hypothesize that the decreased cell viability is a partial consequence of a perturbed cell-cycle progression. In thymocytes, L1/L2-DKO T cells increase the cell cycle progression (15). However, the opposite is true for mature L1/L2-DKO T cells where cell cycle progression is perturbed. Interestingly, L1 or L2 single-deficient T cells did not display a perturbed cell cycle, highlighting the co-regulation of target genes by ZFP36L1 and ZFP36L2. Indeed, Cdkn1a (p21) was already upregulated in the individual absence of ZFP36L1 or ZFP36L2 and Ccne2 (cyclin E2) was only upregulated L1/L2-DKO T cells. How ZFP36L1 and ZFP36L2 co-regulate cell cycle progression in T cells, however, remains to be uncovered.

Mature T cells receive important survival signals through cytokine receptors (32, 33). In line with their reduced cell viability, L1/L2-DKO T cells displayed reduced IL-7R mediated STAT5 signaling. The expression levels of the IL-7R subunits IL7RA and IL2RG were increased in mature L1/L2-DKO T cells. Because IL-7 mediated signaling requires clathrin-mediated endocytosis of IL-7R upon IL-7 binding (34), it is conceivable that internalization of the IL-7R is impaired in L1/L2-DKO T cells. However, while we cannot exclude this possibility, the proteomics analysis did not uncover changes in key components of clathrin-dependent endocytosis, i.e. AP-2 and clathrin in L1/L2-DKO T cells (data not shown). Conversely, the phosphatase DUSP4 known to dephosphorylate STAT5 (31) was induced in L1/L2-DKO T cells, suggesting that the reduced STAT5 phosphorylation is at least in part regulated through this feedback mechanism.

Interestingly, L1/L2-DKO T cells showed increased expression of proteins involved in the unfolded protein response (UPR), including the key reporter of ER-stress EIF2AK3 (PERK) (35). PERK inhibits translation by phosphorylating EIF2A and induces the expression of CHOP through ATF4, which impairs T cell function and induces apoptosis (36). Of note, PERK does not contain AREs in its mRNA. Its induction may therefore be a consequence of increased mRNA stability and thus increased translational demand of L1/L2 target genes during continuous T cell activation, which could potentially lead to the induction of apoptosis via CHOP. If true, this feedback would impede the generation of super-cytokine producers by compromising T cell integrity. This mechanism may thus serve as safeguard to prevent unwanted activity of T cells, and thus to prevent immunopathology, as seen with chronic IFNγ production in hepatitis A infection (37).

In summary, we highlight the important role of ZFP36L1 and ZFP36L2 in coregulating cytokine production during T cell activation, with the most striking effects during continuous activation. In addition, we unveiled the involvement of ZFP36L1 and ZFP36L2 in regulating fundamental cellular processes, including cell cycle progression and cytokine receptor signaling.

## ACKNOWLEDGEMENTS

We thank Dr. Martin Turner for sharing the Zfp36l2;CD4cre (ZFP36L2^cko^) mice; the animal caretakers of the NKI (Netherlands Cancer Institute) and the Sanquin Flow cytometry facility; A. Laurent for technical help; and Koos Rooijers for critical reading of the manuscript. This research was supported by the European Research Council consolidator grant PRINTERS 817533 to M.C.W., and the Landsteiner Foundation of Blood Cell Research 2103 to B.P and M.C.W.

## SUPPLEMENTARY FIGURE LEGENDS

**Supplementary Figure 1. Protein expression kinetics of ZFP36 family members**

**(A**) Immunoblots of ZFP36L2 (upper row), ZFP36L1 (middle row) and ZFP36 (bottom row) protein expression in resting WT or L2-KO (KO) OT-I effector T cells, and T cells activated for indicated time with OVA peptide. RhoGDI levels are used as a loading control. n=2 mice. Presented data is from an individual experiment and one independent experiment has been performed.

**Supplementary Figure 2. Time-dependent regulation of cytokine production by ZFP36L2 and ZFP36L1**

**(A)** Frequency of IL-2 producing OT-I T cells 24h after co-culture with B16-OVA target cells. **(B)** Frequency of TNF producing cells and gMFI levels of TNF (right panel) of TNF producing cells at indicated time points post continuous co-culture with B16-OVA cells. n=3 mice. **(C)** IFNγ producing OT-I T cells and the level of IFNγ production of IFNγ producing OT-I T cells at indicated time points. Brefeldin A was added for the last 2 h of each time point. n=3 mice. (A-C) Presented data is from individual experiments. (A-C) Data of the 24h, 48h and 72h timepoints are representative for three independent experiments. (B-C) Data of the 96h timepoint is from a single experiment. Data were analyzed by one-way ANOVA with Tukey multiple comparison correction, mean ± SD (B, C); *P < 0.05, **P < 0.01, ***P < 0.001, ****P < 0.0001.

**Supplementary Figure 3. Characterization and validation of differentially expressed proteins between WT and L1/L2-DKO T cells**

**(A)** Protein expression levels of ZFP36L2, ZFP36L1 and ZFP36 in log2 intensity, as determined by mass spectrometry (MS) analysis. n=2-5 mice per condition. **(B)** Significantly enriched regions (P.adj< 0.05) bound by ZFP36L1 identified using iCLIP data. Data for analysis was retrieved from (14). n=3 mice per genotype. **(C)** Number of total target (from B) transcripts bound by ZFP36L1, and regions of ZFP36L1 interactions within different regions. **(D)** ZFP36L1 CLIP data on the 3’UTR region of *Tnf*, *Ifng* and *IL-2* mRNA. Red bar: AU-rich elements. Each lane depicts CLIP data of ZFP36L1-proficient (WT) or ZFP36L1-deficient (KO) T cells activated for 3h with OVA peptide. n=3 mice per genotype. **(E)** Venn diagram depicting the overlap of transcripts from ZFP36L1 CLIP data with higher intensity values (log2 fold change > 0.5) in L1/L2-DKO T cells during resting and 24h of activation. **(F)** Dot plots of log2 intensity of expression of selected proteins with higher intensities (log2 fold change > 0.5) in L1/L2-DKO T cells, and that were identified in ZFP36L1 CLIP data. Each dot indicates one mouse. n=2-5 mice. **(G)** Heatmap of the relative expression levels of proteins with different intensities (P.adj < 0.05 & Log2FC > or < 0.5) between WT and L1/L2-DKO T cells, associated with the unfolded protein response. **(H)** Dot plots showing the protein expression levels in log2 intensity for Ifngr1. **(I-J)** Heatmap of the relative expression levels of proteins with different intensities (P.adj < 0.05 & Log2FC > or < 0.5) between WT and L1/L2-DKO T cells, associated with the G2M checkpoint (I) and E2F targets (J) based on Hallmark Gene Set Enrichment Analysis (GSEA; Fig. 4D). n=2-5 mice. (A-J) Presented data is from individual experiments. (A-J) One independent experiment has been performed. (A, F-I): Asteriks indicate significantly differentially abundant proteins between WT and L1/L2- DKO cells; P.adj < 0.05 & Log2FC > or < 0.5.

**Supplementary Figure 4. Dysregulated cell cycle progression and IL-7 stat5 mediated signaling in L1/L2-DKO T cells**

**(A)** Relative ratio of dying (LiveDead^int^) OT-I cells present in G0, G1 and S-M cell cycle. n=3 mice. **(B-C)** Cell cycle analysis of OT-I T cells using Ki-67 and PI after 4h (G) and 24h (H) of activation with 100nM OVA peptide. n=3 mice. **(D)** Histograms showing Cell Trace Yellow (CTY) levels in OT-I T cells that were cultured in the absence (left) or presence of 100nM OVA peptide for 48h. Relative CTY signal among 48h activated samples is calculated by using the 48h unactivated T cells as a reference. n=3 mice. Data were analyzed by one-way ANOVA with Tukey multiple comparison correction, mean ± SD (A, B, C, D); *P < 0.05, **P < 0.01, ****P < 0.0001. **(E)** Heatmap showing relative protein expression levels of differentially expressed proteins (P.adj < 0.05 & Log2FC > or < 0.5) that are associated with the IFNγ response in resting cells (Fig. 4D). **(F)** Dot plots showing the protein expression levels in log2 intensity for Ifngr1. **(G)** Heatmap showing relative protein expression levels of differentially expressed proteins (P.adj < 0.05 & Log2FC > or < 0.5) that are associated with the IL2 stat5 signalling in resting cells (Fig. 4D). **(H-I)** Log2 intensity of protein expression levels for Stat5a and Stat5b (H), and Dusp4 (I) n=2-5 mice. (A-I) Presented data is from individual experiments. (A) Data is representative for three independent experiments. (B-I) One independent experiment has been performed. Asteriks indicates significantly differentially expressed proteins between WT and L1/L2-DKO cells; P.adj < 0.05 & Log2FC > or < 0.5.

## MATERIALS AND METHODS

### Mice and cell culture

OTI;Zfp36l2;CD4cre and C57BL/6j/Ly5.1^+^Ly5.2^+^ mice were bred and housed in the animal facility of the Netherlands Cancer Institute (NKI). OTI;Zfp36l2;CD4cre^negative^ mice were used to obtain WT OT-I T cells and OTI;Zfp36l2;CD4cre^positive^ mice were used to obtain ZFP36L2-KO OT-I T cells (L2-KO). Animals were genotyped by Transnetyx. Animals were housed in individually ventilated cage systems under specific pathogen-free conditions. All experiments were performed in accordance with institutional and national guidelines and approved by the experimental animal committee of the NKI. Mice were used at 6-12 weeks of age. B16-OVA cells (24) and MEC.B7.SigOVA cells (38) were cultured at 37°C and 5% CO_2_ in IMDM (GIBCO-BRL) containing L-glutamine and 25mM HEPES, supplemented with 10% FCS, 15μM 2-mercaptopethanol, 2mM L-glutamine, 100 U/ml penicillin G sodium and 100 μg/ml streptomycin sulfate.

### T cell isolation and activation

Mouse spleens, inguinal and mesenteric lymph nodes were harvested and briefly kept in cold PBS on ice before being mashed and passed through a 70μm cell strainer (Corning) and washed with FCS-containing media. Erythrocytes were lysed using cold red blood cell lysis buffer (155 mM NH4Cl, 10 mM KHCO3, 0.1 mM EDTA, pH 7.4) for 60s. OT-I T cells were purified with the MACS CD8a^+^ T cell isolation kit (Miltenyi Biotec) according to the manufacturer’s protocol. To generate effector OT-I T cells, purified OT-I T cells (1e6 cells/well) were activated with pre-seeded MEC.B7.SigOVA cells (1e5 cells/well) for 20h. T cells were removed from stimulation and cultured in complete medium supplemented with 10 ng/ml recombinant mouse IL-7 (Peprotech) for 3-5 days as previously described (39). In vitro reactivation of OT-I T cells was performed with 100μM of OVA peptide_257-264_. For cytokine measurements, T cells were incubated with 1μg/ml Brefeldin A (BD Biosciences) for the last 2h of activation.

### Genetic modification of primary mouse T cells with Cas9-RNPs

ZFP36L1 crRNAs were designed and mouse T cell nucleofection was performed as previously described (18). Briefly, ALT-R crRNA and tracr-RNA were reconstituted to 100uM in Nuclease Free Duplex buffer (all Intergrated DNA Technologies). Non-targeting negative control crRNA #1 (Integrated DNA Technologies) were used as negative control. crRNA and tracrRNA were mixed in equimolar ratios (4.5μl total crRNA + 4.5μl tracrRNA) in nuclease-free PCR tubes and denatured at 95°C for 5m in a thermocycler and cooled down to room temperature prior to adding 30mg of SpCas9 (Integrated DNA Technologies) to generate ribonuclear proteins (RNPs). OT-I T cells activated for 20h with MEC.B7.SigOVA cells and rested for an additional 24h in medium containing IL-7 (10ng/ml) were washed with PBS, resuspended in P2 buffer (Lonza) combined with RNPs, and electroporated in a 16-well 4D Nucleofector X strip (Lonza) using the CM137 program. ZFP36L1 knockout efficiency was determined 3 days after electroporation by Western blot.

### Western blotting

For Western blotting, OT-I T cells were washed with PBS (1ml), pelleted, snap-frozen with liquid nitrogen, and stored at −80°C until further use. Cell pellets were lysed using RIPA buffer (ThermoFisher) supplemented with 10mM DTT and Halt Protease and Phosphatase inhibitors (100X, ThermoFisher). Samples were run using Laemmli SDS sample buffer (ThermoFisher) on 4-12% SDS-PAGE gels (Invitrogen). Proteins were transferred onto a nitrocellulose membrane using the iBlot2 system (ThermoFisher). Nitrocellulose membranes were blocked for 1h in TBST buffer containing 5% BSA (Sigma) prior to o/n staining with antibodies targeting ZFP36L2 (ab70775, Abcam), ZFP36L1 (ab42473, Abcam), ZFP36 (#OTI3D10, Origene) and RhoGDI (MAB9959, Abnova). Secondary antibody staining was performed with goat a-rabbit (4050–05) or goat a-mouse-HRP (1031-05, both Southern Biotech).

### B16-OVA tumor model

C57BL/6J/Ly5.1^+^Ly5.2^+^ mice (8-12-week-old) were injected with 1 x 10^6^ B16-OVA cells. On day 7, when tumors were palpable, *in vitro* generated WT, L1-KO, L2-KO, or L1-L2 DKO OT-I T cells that were rested and expanded in rIL-7 for 3 days after nucleofection were washed twice with PBS. 1 x 10^6^ cells were administered intravenously via retro-orbital injection. At day 16 post tumor injection, animals were sacrificed and tumors were harvested.

Tumors were manually dissociated using scalpel blades and digested using Collagenase IV (200 U/ml, Worthington Biochemical) and DNase I (100 μg/ml, Roche) for 30m at 37°C. Cells were incubated with 1μg/ml brefeldin A and 1μg/ml monensin for 4h in the presence or absence of 100nM OVA_257-264_ peptide.

For *in vitro* B16-OVA co-culture experiments, OT-I T cells rested for 4-5 days of indicated genetic background were co-cultured with B16-OVA cells that were seeded 3h prior to start of co-culture at a 6:1 effector:target ratio for 6h or 24h in a 24-well plate. For timepoints longer than 24h, T cells were transferred each day to freshly seeded B16-OVA cells.

### Quantitative RT-PCR analysis

Total RNA was extracted from OT-I T cells using Trizol reagent (Invitrogen). cDNA was synthesized using Superscript III Reverse transcriptase (Invitrogen). Quantitative Real-Time PCR was performed in triplicate with SYBR green on a StepOne Plus (both Applied Biosystems) using previously described primers for *Ifng*, *Tnf*, *IL-2* and *Rpl32* mRNA (39). Ct values were normalized to RPL32 levels. *Ifng* mRNA decay was determined upon treatment with 1 μg/ml of Actinomycin D (Sigma-Aldrich) for indicated time points.

### Flow cytometry

Cells were washed with cold FACS buffer (PBS, 1% FCS, and 2mM EDTA) and stained with LIVE/DEAD Near-IR Dead cell dye (Invitrogen), according to the manufacturer’s protocol. For extracellular stainings, anti-CD8a (53-6.7, BD) and anti-CD127 (A7R34, Biolegend) were incubated for 20 min at 4°C. For intracellular cytokine staining, cells were fixed and permeabilized using the BD Cytofix/Cytoperm kit (BD Biosciences) according to the manufacturer’s protocol. Cytokines were stained with the following antibodies: anti-TNF (MP6-XT22, Biolegend), anti-IFN-γ (XMG1.2, Biolegend), anti-IL-2 (JES6-5H4, ThermoFisher Scientific). For cell cycle analysis, cells were fixed using the Foxp3/transcription factor staining kit (ThermoFisher Scientific) according to the manufacturer’s protocol and stained with Ki-67 (SolA15, eBioscience). For the staining of phosphorylated STAT5 (pSTAT5), after 20m exposure to IL-7 (15ng/ml), cells were washed with cold FACS buffer. Prior to staining, cells were fixed using the Foxp3/transcription factor staining kit (ThermoFisher Scientific) followed by fixation with 90% methanol for 15 min. Cells were subsequently stained overnight in FACS buffer with anti-pSTAT5 (SRBCZX, ThermoFisher Scientific). Flow cytometry analysis was performed with FACSymphony (BD Biosciences) and data were analyzed with FlowJo (BD Biosciences, version 10).

### IFN**γ** blocking experiment

24h after nucleofection, IFNγ blocking antibody (XMG1.2, Invitrogen) was added to the culture at 10μg/ml every 48h. 5 days after nucleofection, the viability of cells was measured using the LIVE/DEAD Near-IR Dead cell dye (Invitrogen).

### Cell cycle and proliferation analysis

Cell cycle analysis was performed on T cells rested for 4 days in the presence of rIL-7. After extracellular and intracellular staining, cells were washed with FACS buffer and stained for DNA content using Propidium Iodide (Invitrogen) in the presence of 10μg/ml RNAse (Qiagen). For proliferation assays, cells were washed with PBS and stained the CellTrace Yellow proliferation dye (Invitrogen) according to the manufacturer’s protocol.

### Mass spectrometry sample preparation

1 x 10^6^ WT, L2-KO and L1-KO, and 2 x 10^6^ L1/L2-DKO resting T cells were left untreated or activated with OVA257-264 peptide for a period of 24h. Cells were pelleted and washed twice with ice-cold PBS. Cell pellets were snap frozen using liquid nitrogen and stored at - 70°C until further processing. Cell pellets were lysed in 1% sodium deoxycholate (SDC, BioWORLD, Dublin, OH, USA), 10 mM Tris-(2-carboxyethyl)phosphine (Thermo Fisher Scientific, Rockford, IL, USA), 40 mM Chloroacetamide (Sigma-Aldrich, Saint Louis, MO, USA), 100 mM Tris pH 8, (Life Technologies, Thermo Fisher Scientific, Rockford, IL, USA) and heated for 5 min at 95 °C. After cooling to room temperature, the samples were sonicated in a sonication waterbath (Branson Ultrasonics, Brookfield, CT, USA) for 10 min. Proteins were digested for 18 h at 25 °C with trypsin/LysC (Thermo Fisher Scientific, Rockford, IL, USA). Peptides were separated on an Evosep one (Evosep, Odense, Denmark) with the preset 30 samples-per-day method on a 15 cm Evosep Performance Column (EV-1137, 150 µm I.D., 1.5 µm particle size). Acquisition was performed on a timsTOF-HT (Bruker Daltonics, Billerica, MA, USA) mass spectrometer operated in dia-PASEF mode. Ion mobility accumulation and ramp time were set to 100 ms. Further, 16 DIA windows were set per cycle, ranging from 0.7–1.5 1/k0 to 421–1594 m/z, and the size of DIA windows was set based on precursor density. Collision energy was set as a linear function of the ion mobility (0.6 1/k0 = 20 eV, 1.60 1/k0 = 59 eV). Raw files were processed in DIA-NN 1.8.1.; proteins and peptides were detected by querying the filtered mouse Swissprot database (release 28-08-2024). Standard settings were used, using a generated library-based spectra search. Maximum number of variable modifications was set to 2. Protein Interference used was “Protein names (from FASTA)” and quantification strategy “Robust LC (high precision)”. Data were analyzed using R 4.4.1/Rstudio (2024.9.1.394). Detected proteins were filtered for proteotypic and ≥2 unique peptides per protein, and proteins were quantified in 100% of samples in at least one condition.Proteins quantified in at least one experimental group were selected for further analysis. Missing values were imputed by using a down-shifted normal distribution (width=0.3, shift = 1.8) according to Tyanova *et al.* (40), assuming these proteins were close to the detection limit. Statistical analyses were performed between WT and L1/L2-DKO samples using moderated t-tests in the LIMMA package. A Benjamini-Hochberg adjusted P value <0.05 and absolute log2fold change >0.5 was considered statistically significant. Proteins that met the log2 fold change threshold were taken along for visualization. K-means clustering was performed using the stats package that is part of R. Gene Set Enrichment Analysis was performed using the mouse-ortholog hallmark gene sets from the Molecular Signatures Database (41, 42). The R package ggplot2 was used for graphical representations.

### iCLIP analysis

iCLIP dataset for ZFP36L1 in CD8^+^ T cells following 3h of activation with 0.1μM OVA peptide were obtained from GSE176313 (14). Adaptor sequences were removed from sequence reads using Cutadapt (43). Sequencing reads were mapped to the GRCm39 mouse genome using STAR (44) and deduplicated using the random barcodes with Samtools (45). ZFP36L1 crosslink sites were extracted and counted using HTSeq (46). Significant differentially enriched regions (adjusted P-value <0.05) were further identified using DEWSeq (47).

### Statistical analysis

Data is presented as mean ± standard deviation. Statistical analysis was performed using GraphPad Prism 10. Statistical analysis was performed using the mean and standard deviation values. A paired two-sided student *t*-test was used when comparing two groups, while an ordinary one-way ANOVA test with a Tukey correction was used when comparing more than two groups. P values < 0.05 were considered.

## Notes

### Competing Interest Statement

The authors have declared no competing interest.

## REFERENCES

1. Zhang, B., T. Karrison, D. A. Rowley, and H. Schreiber. 2008. IFN-γ– and TNF-dependent bystander eradication of antigen-loss variants in established mouse cancers. J. Clin. Invest. 118: 1398–1404.

2. Ciuffreda, D., D. Comte, M. Cavassini, E. Giostra, L. Bühler, M. Perruchoud, M. H. Heim, M. Battegay, D. Genné, B. Mulhaupt, R. Malinverni, C. Oneta, E. Bernasconi, M. Monnat, A. Cerny, C. Chuard, J. Borovicka, G. Mentha, M. Pascual, J.-J. Gonvers, G. Pantaleo, and V. Dutoit. 2008. Polyfunctional HCV-specific T-cell responses are associated with effective control of HCV replication. Eur. J. Immunol. 38: 2665–2677.

3. Boulch, M., M. Cazaux, A. Cuffel, M. V. Guerin, Z. Garcia, R. Alonso, F. Lemaître, A. Beer, B. Corre, L. Menger, C. L. Grandjean, F. Morin, C. Thieblemont, S. Caillat-Zucman, and P. Bousso. 2023. Tumor-intrinsic sensitivity to the pro-apoptotic effects of IFN-γ is a major determinant of CD4+ CAR T-cell antitumor activity. Nat Cancer.

4. Grivennikov, S. I., A. V. Tumanov, D. J. Liepinsh, A. A. Kruglov, B. I. Marakusha, A. N. Shakhov, T. Murakami, L. N. Drutskaya, I. Förster, B. E. Clausen, L. Tessarollo, B. Ryffel, D. V. Kuprash, and S. A. Nedospasov. 2005. Distinct and Nonredundant In Vivo Functions of TNF Produced by T Cells and Macrophages/Neutrophils. Immunity 22: 93–104.

5. Salerno, F., S. Engels, M. van den Biggelaar, F. P. J. van Alphen, A. Guislain, W. Zhao, D. L. Hodge, S. E. Bell, J. P. Medema, M. von Lindern, M. Turner, H. A. Young, and M. C. Wolkers. 2018. Translational repression of pre-formed cytokine-encoding mRNA prevents chronic activation of memory T cells. Nature Immunology 19: 828–837.

6. Salerno, F., N. A. Paolini, R. Stark, M. von Lindern, and M. C. Wolkers. 2017. Distinct PKC-mediated posttranscriptional events set cytokine production kinetics in CD8 ^+^ T cells. Proceedings of the National Academy of Sciences 114: 201704227.

7. Han, J., Y. Zhao, K. Shirai, A. Molodtsov, F. W. Kolling, J. L. Fisher, P. Zhang, S. Yan, T. G. Searles, J. M. Bader, J. Gui, C. Cheng, M. S. Ernstoff, M. J. Turk, and C. V. Angeles. 2021. Resident and circulating memory T cells persist for years in melanoma patients with durable responses to immunotherapy. Nat Cancer 2: 300–311.

8. Veiga-Fernandes, H., U. Walter, C. Bourgeois, A. McLean, and B. Rocha. 2000. Response of naïve and memory CD8+ T cells to antigen stimulation in vivo. Nat Immunol 1: 47–53.

9. Fara, A., Z. Mitrev, R. A. Rosalia, and B. M. Assas. 2020. Cytokine storm and COVID-19: a chronicle of pro-inflammatory cytokines. Open Biol. 10: 200160.

10. Morgan, R. A., J. C. Yang, M. Kitano, M. E. Dudley, C. M. Laurencot, and S. A. Rosenberg. 2010. Case Report of a Serious Adverse Event Following the Administration of T Cells Transduced With a Chimeric Antigen Receptor Recognizing ERBB2. Molecular Therapy 18: 843–851.

11. Jurgens, A. P., B. Popović, and M. C. Wolkers. 2021. T cells at work: How post-transcriptional mechanisms control T cell homeostasis and activation. European Journal of Immunology 51: 2178–2187.

12. Nicolet, B. P., N. D. Zandhuis, V. M. Lattanzio, and M. C. Wolkers. 2021. Sequence determinants as key regulators in gene expression of T cells. Immunological reviews 1–20.

13. Anderson, P. 2008. Post-transcriptional control of cytokine production. Nat Immunol 9: 353–359.

14. Petkau, G., T. J. Mitchell, K. Chakraborty, S. E. Bell, V. D’Angeli, L. Matheson, D. J. Turner, A. Saveliev, O. Gizlenci, F. Salerno, P. D. Katsikis, and M. Turner. 2022. The timing of differentiation and potency of CD8 effector function is set by RNA binding proteins. Nature Communications 13.

15. Galloway, A., H. Ahlfors, M. Turner, L. S. Bell, and K. U. Vogel. 2016. The RNA-Binding Proteins Zfp36l1 and Zfp36l2 Enforce the Thymic β-Selection Checkpoint by Limiting DNA Damage Response Signaling and Cell Cycle Progression. The Journal of Immunology 197: 2673–2685.

16. Cook, M. E., T. R. Bradstreet, A. M. Webber, J. Kim, A. Santeford, K. M. Harris, M. K. Murphy, J. Tran, N. M. Abdalla, E. A. Schwarzkopf, S. C. Greco, C. M. Halabi, R. S. Apte, P. J. Blackshear, and B. T. Edelson. 2022. The ZFP36 family of RNA binding proteins regulates homeostatic and autoreactive T cell responses. Science Immunology 0981.

17. Petkau, G., T. J. Mitchell, M. J. Evans, L. Matheson, F. Salerno, and M. Turner. 2023. Zfp36l1 establishes the high[:affinity CD8 T[:cell response by directly linking TCR affinity to cytokine sensing. Eur J Immunol 2350700.

18. Popović, B., B. P. Nicolet, A. Guislain, S. Engels, A. P. Jurgens, N. Paravinja, J. J. Freen-van Heeren, F. P. J. Van Alphen, M. Van Den Biggelaar, F. Salerno, and M. C. Wolkers. 2023. Time-dependent regulation of cytokine production by RNA binding proteins defines T cell effector function. Cell Reports 42: 112419.

19. Zandhuis, N. D., A. Guislain, A. Popalzij, S. Engels, B. Popović, M. Turner, and M. C. Wolkers. 2024. Regulation of IFN[:γ production by ZFP36L2 in T cells is time[:dependent. Eur J Immunol 2451018.

20. Caput, D., B. Beutler, K. Hartog, R. Thayer, S. Brown-Shimer, and A. Cerami. 1986. Identification of a common nucleotide sequence in the 3’-untranslated region of mRNA molecules specifying inflammatory mediators. Proceedings of the National Academy of Sciences of the United States of America 83: 1670–1674.

21. Matheson, L. S., G. Petkau, B. Sáenz-Narciso, V. D’Angeli, J. McHugh, R. Newman, H. Munford, J. West, K. Chakraborty, J. Roberts, S. Łukasiak, M. D. Díaz-Muñoz, S. E. Bell, S. Dimeloe, and M. Turner. 2022. Multiomics analysis couples mRNA turnover and translational control of glutamine metabolism to the differentiation of the activated CD4+ T cell. Sci Rep 12: 19657.

22. Moore, M. J., N. E. Blachere, J. J. Fak, C. Y. Park, K. Sawicka, S. Parveen, I. Zucker-Scharff, B. Moltedo, A. Y. Rudensky, and R. B. Darnell. 2018. ZFP36 RNA-binding proteins restrain T cell activation and anti-viral immunity. eLife 7: 920–926.

23. Brenes, A. J., A. I. Lamond, and D. A. Cantrell. 2023. The Immunological Proteome Resource. Nat Immunol 24: 731–731.

24. De Witte, M. A., M. Coccoris, M. C. Wolkers, M. D. Van Den Boom, E. M. Mesman, J.-Y. Song, M. Van Der Valk, J. B. A. G. Haanen, and T. N. M. Schumacher. 2006. Targeting self-antigens through allogeneic TCR gene transfer. Blood 108: 870–877.

25. Salerno, F., A. Guislain, J. J. Freen-Van Heeren, B. P. Nicolet, H. A. Young, and M. C. Wolkers. 2019. Critical role of post-transcriptional regulation for IFN-γ in tumor-infiltrating T cells. OncoImmunology 8: 1–12.

26. Dighe, A. S., E. Richards, L. J. Old, and R. D. Schreiber. 1994. Enhanced in vivo growth and resistance to rejection of tumor cells expressing dominant negative IFNγ receptors. Immunity 1: 447–456.

27. Bakheet, T., E. Hitti, M. Al-Saif, W. N. Moghrabi, and K. S. A. Khabar. 2018. The AU-rich element landscape across human transcriptome reveals a large proportion in introns and regulation by ELAVL1/HuR. Biochimica et Biophysica Acta - Gene Regulatory Mechanisms 1861: 167–177.

28. Miller, I., M. Min, C. Yang, C. Tian, S. Gookin, D. Carter, and S. L. Spencer. 2018. Ki67 is a Graded Rather than a Binary Marker of Proliferation versus Quiescence. Cell Reports 24: 1105–1112.e5.

29. Refaeli, Y., L. Van Parijs, S. I. Alexander, and A. K. Abbas. 2002. Interferon γ Is Required for Activation-induced Death of T Lymphocytes. Journal of Experimental Medicine 196: 999–1005.

30. Vella, A. T., S. Dow, T. A. Potter, J. Kappler, and P. Marrack. 1998. Cytokine-induced survival of activated T cells *in vitro* and *in vivo*. Proc. Natl. Acad. Sci. U.S.A. 95: 3810–3815.

31. Hsiao, W.-Y., Y.-C. Lin, F.-H. Liao, Y.-C. Chan, and C.-Y. Huang. 2015. Dual-Specificity Phosphatase 4 Regulates STAT5 Protein Stability and Helper T Cell Polarization*. PLoS ONE 10: e0145880.

32. Vella, A. T., S. Dow, T. A. Potter, J. Kappler, and P. Marrack. 1998. Cytokine-induced survival of activated T cells *in vitro* and *in vivo*. Proc. Natl. Acad. Sci. U.S.A. 95: 3810–3815.

33. Murray, R., T. Suda, N. Wrighton, F. Lee, and A. Ziotnik. 1989. IL-7 is a growth and maintenance factor for mature and immature thymocyte subsets. Int Immunol 1: 526–531.

34. Henriques, C. M., J. Rino, R. J. Nibbs, G. J. Graham, and J. T. Barata. 2010. IL-7 induces rapid clathrin-mediated internalization and JAK3-dependent degradation of IL-7Rα in T cells. Blood 115: 3269–3277.

35. Harding, H. P., Y. Zhang, and D. Ron. 1999. Protein translation and folding are coupled by an endoplasmic-reticulum-resident kinase. Nature 397: 271–274.

36. Cao, Y., J. Trillo-Tinoco, R. A. Sierra, C. Anadon, W. Dai, E. Mohamed, L. Cen, T. L. Costich, A. Magliocco, D. Marchion, R. Klar, S. Michel, F. Jaschinski, R. R. Reich, S. Mehrotra, J. R. Cubillos-Ruiz, D. H. Munn, J. R. Conejo-Garcia, and P. C. Rodriguez. 2019. ER stress-induced mediator C/EBP homologous protein thwarts effector T cell activity in tumors through T-bet repression. Nat Commun 10: 1280.

37. Kim, J., D.-Y. Chang, H. W. Lee, H. Lee, J. H. Kim, P. S. Sung, K. H. Kim, S.-H. Hong, W. Kang, J. Lee, S. Y. Shin, H. T. Yu, S. You, Y. S. Choi, I. Oh, D. H. Lee, D. H. Lee, M. K. Jung, K.-S. Suh, S. Hwang, W. Kim, S.-H. Park, H. J. Kim, and E.-C. Shin. 2018. Innate-like Cytotoxic Function of Bystander-Activated CD8+ T Cells Is Associated with Liver Injury in Acute Hepatitis A. Immunity 48: 161–173.e5.

38. Van Stipdonk, M. J. B., E. E. Lemmens, and S. P. Schoenberger. 2001. Naïve CTLs require a single brief period of antigenic stimulation for clonal expansion and differentiation. Nat Immunol 2: 423–429.

39. Salerno, F., A. Guislain, D. Cansever, and M. C. Wolkers. 2016. TLR-Mediated Innate Production of IFN-γ by CD8 + T Cells Is Independent of Glycolysis. The Journal of Immunology 196: 3695–3705.

40. Tyanova, S., T. Temu, P. Sinitcyn, A. Carlson, M. Y. Hein, T. Geiger, M. Mann, and J. Cox. 2016. The Perseus computational platform for comprehensive analysis of (prote)omics data. Nat Methods 13: 731–740.

41. Subramanian, A., P. Tamayo, V. K. Mootha, S. Mukherjee, B. L. Ebert, M. A. Gillette, A. Paulovich, S. L. Pomeroy, T. R. Golub, E. S. Lander, and J. P. Mesirov. 2005. Gene set enrichment analysis: A knowledge-based approach for interpreting genome-wide expression profiles. Proc. Natl. Acad. Sci. U.S.A. 102: 15545–15550.

42. Castanza, A. S., J. M. Recla, D. Eby, H. Thorvaldsdóttir, C. J. Bult, and J. P. Mesirov. 2023. Extending support for mouse data in the Molecular Signatures Database (MSigDB). Nat Methods 20: 1619–1620.

43. Martin, M. 2011. Cutadapt removes adapter sequences from high-throughput sequencing reads. EMBnet j. 17: 10.

44. Dobin, A., C. A. Davis, F. Schlesinger, J. Drenkow, C. Zaleski, S. Jha, P. Batut, M. Chaisson, and T. R. Gingeras. 2013. STAR: ultrafast universal RNA-seq aligner. Bioinformatics 29: 15–21.

45. Danecek, P., J. K. Bonfield, J. Liddle, J. Marshall, V. Ohan, M. O. Pollard, A. Whitwham, T. Keane, S. A. McCarthy, R. M. Davies, and H. Li. 2021. Twelve years of SAMtools and BCFtools. GigaScience 10: giab008.

46. Anders, S., P. T. Pyl, and W. Huber. 2015. HTSeq—a Python framework to work with high-throughput sequencing data. Bioinformatics 31: 166–169.

47. Schwarzl, T., S. Sahadevan, B. Lang, M. Miladi, R. Backofen, W. Huber, M. W. Hentze, and G. G. Tartaglia. 2024. Improved discovery of RNA-binding protein binding sites in eCLIP data using DEWSeq. Nucleic Acids Research 52: e1–e1.

