## Supplementary figures and images for "Combined deletion of ZFP36L1 and ZFP36L2 drives superior cytokine production in T cells at the cost of cell fitness"

### Suppl. Fig 1

#### Supplementary Figure 1

## A

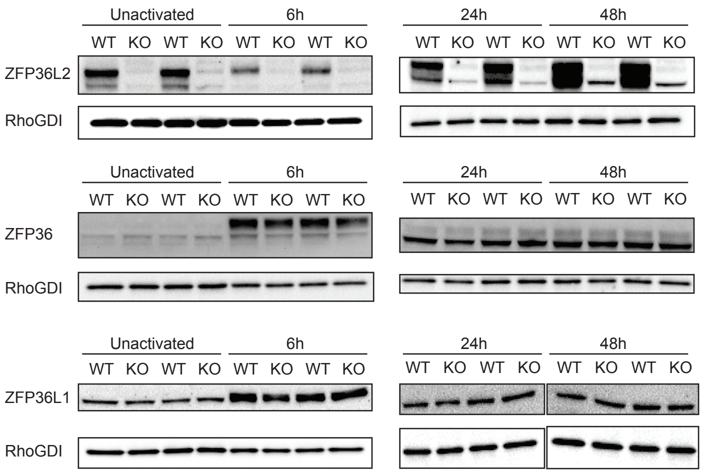

### Suppl. Fig 2

#### Supplementary Figure 2

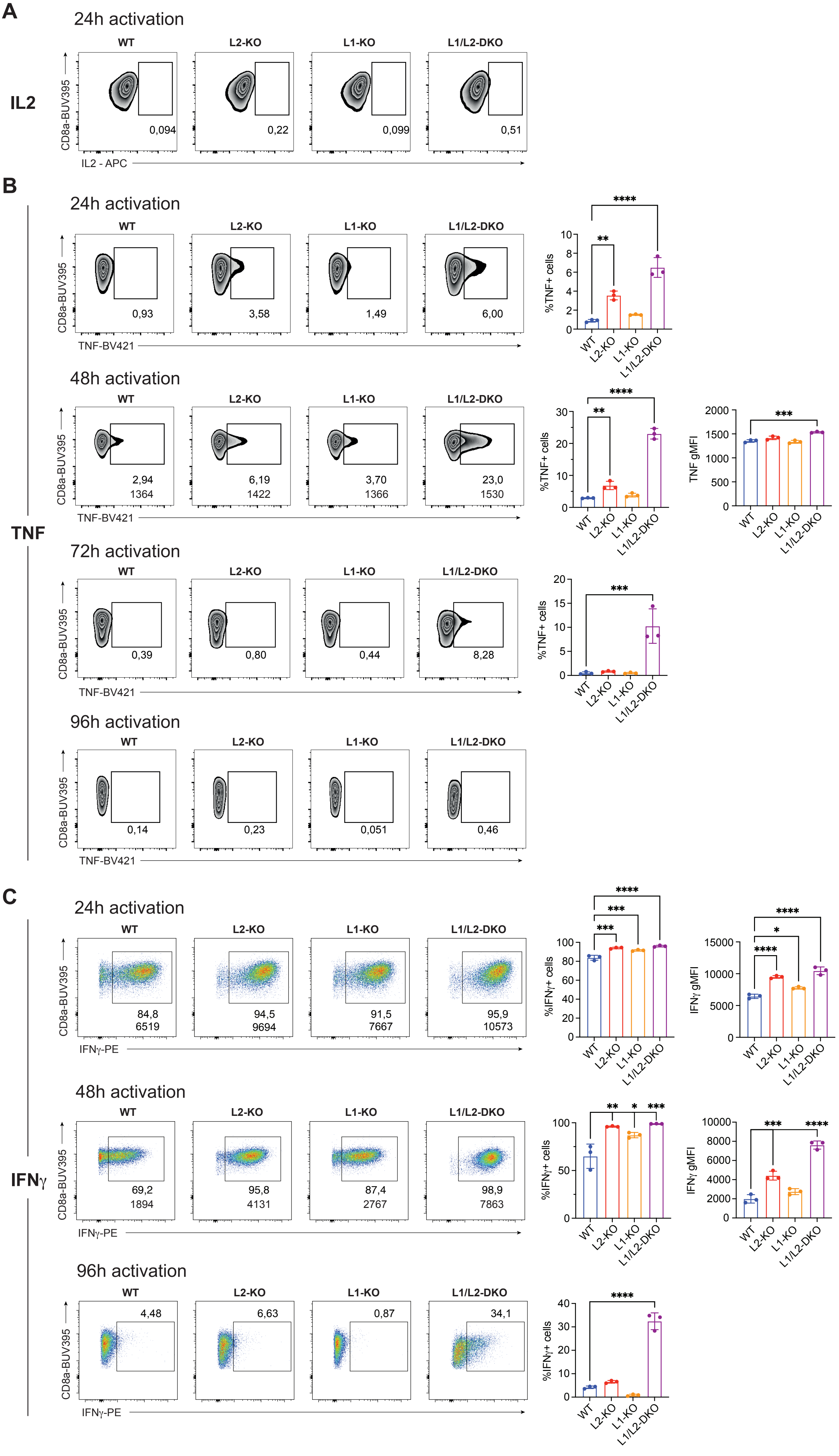

### Suppl. Fig 3

#### Supp. Figure 3

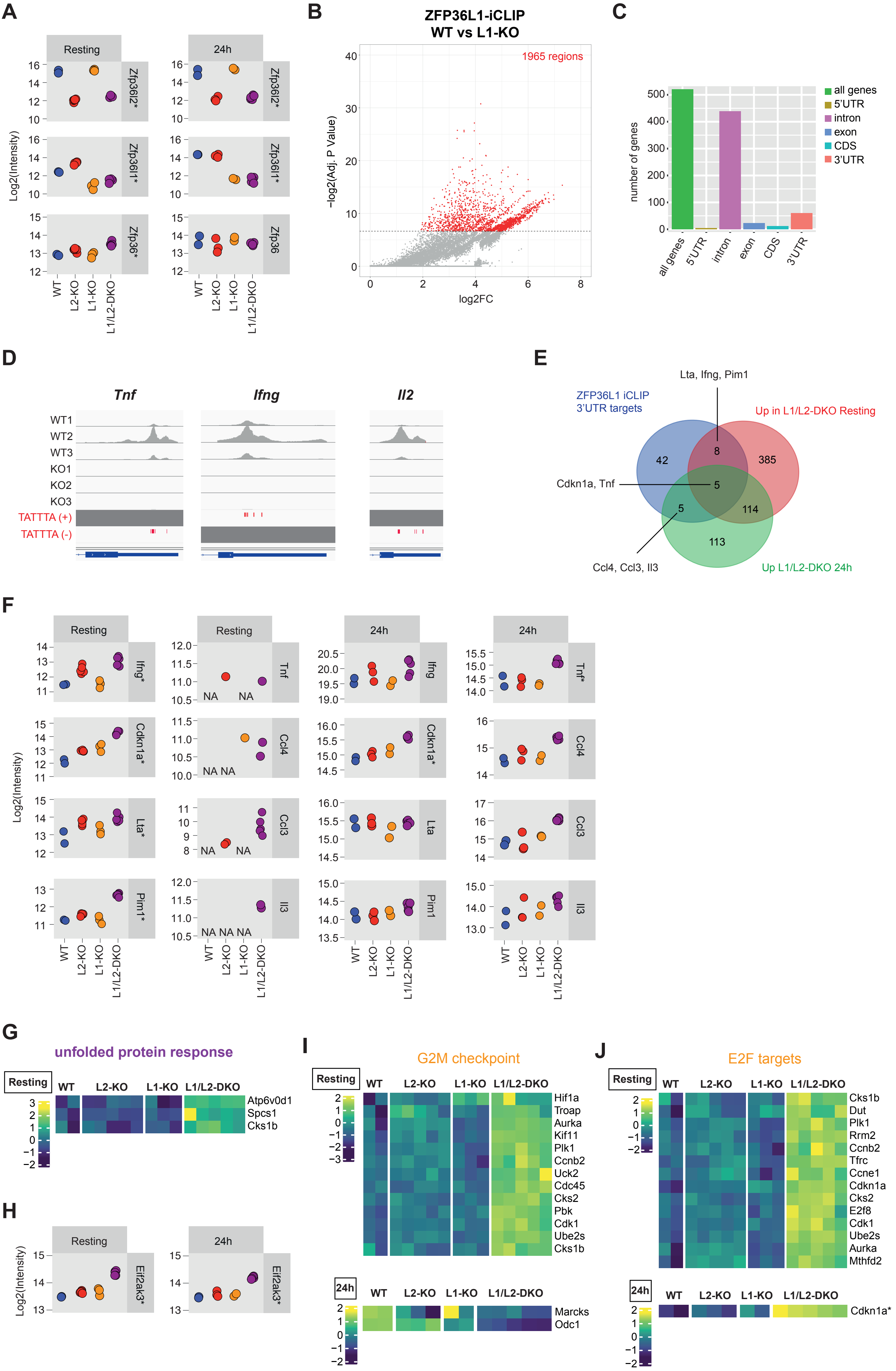

### Suppl. Fig 4

#### Supplementary Figure 4

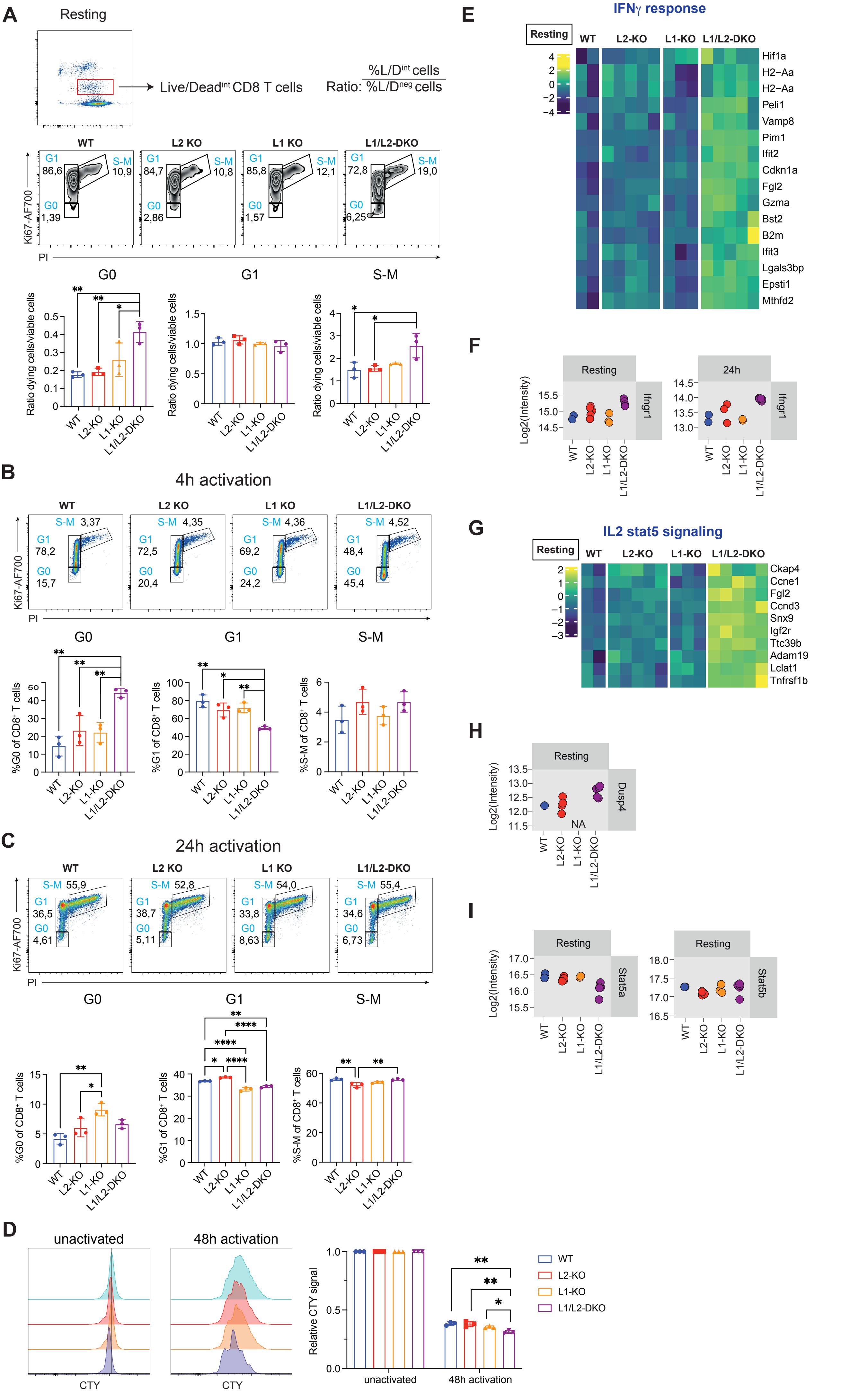
